# Non-Contrast µCT Analysis of Obese Adipose in Response to Cold-Exposure Reveals Sex-Specific Alterations

**DOI:** 10.1101/2025.01.04.631266

**Authors:** Austin E. Eades, Macy M. Payne, Zachary Clark, Logan Evans, R. McKinnon Walsh, Bailey B. Bye, Michaella J. Rekowski, E. Matthew Morris, Stefan H. Bossmann, Michael N. VanSaun

## Abstract

In 2020, it was reported that obesity in the United States had increased by 12% from 1999 to 2018. While exercise and diet are optimal lifestyle modifications to curb obesity, drug-based therapeutics focus on glucagon-like peptide (GLP) modifiers. Alternatively, current research suggests that a specialized type of adipose, called thermogenic adipose, may help protect against obesity. Active thermogenic adipose can metabolize free fatty acids (FFAs) and carbohydrates to carry out non-shivering thermogenesis (NST), potentially providing a method for reducing excess energy stores. While brown adipose tissue (BAT) provides the primary thermogenic response, we hypothesized that exposing diet-induced obese (DIO) mice to colder temperatures would also diminish white adipose tissue (WAT) depots and suppress their inflammatory signature. To measure adipose response to cold *in vivo*, we applied a non-contrast microCT (µCT) imaging analysis. Male and female mice were housed at thermoneutrality (TN) and fed a Western-style diet (WD) *ad lib* until they became obese. Once they reached this stage, the mice were subjected to a thermalshift (TS) and exposed to either room temperature (RT) of 22°C or a colder temperature of 18°C. The adipose response was then assessed in post-exposure tissues by histological analysis, proteomics, and molecular characterization to correlate phenotypic changes with our µCT findings. Results from this analysis revealed a sex-specific response to cold exposure: thermogenic adipose was predominantly formed in the interscapular BAT (iBAT) of male mice, while female mice showed formation in their perigonadal WAT (pgWAT) and iBAT when exposed to 18°C. Furthermore, male mice exhibited a decline in serum glucose levels when subjected to 18°C, which was increased in TS female mice. Serum-free fatty acids (FFAs) were unaffected by either sex across different environmental conditions. Importantly, using a mass-spectrometry-based approach, we detected a reduction in pro-inflammatory cytokines in the conditioned media (CM) of pgWAT and iBAT from TS male mice compared to TN DIO male mice. Overall, our studies demonstrated a new µCT-based analytical method to detect changes in obese adipose tissue and highlighted unique sex-specific responses to environmental exposure. Our findings suggest that thermogenic adipose may offer a promising avenue for combating obesity and reducing its pathologic characteristics.

## Introduction

Obesity has become a significant health crisis in the United States and around the world. [1, 2]. Individuals with obesity have a greater prevalence for the development of metabolic diseases (e.g., hyperglycemia, hyperinsulinemia, hypertension) and low-grade inflammation [1, 3–5]. A significant source of inflammatory cytokines is the white adipose tissue (WAT), which primarily results from adipocyte hypertrophy with increasing BMI. Strategies have been developed to reduce the adipocyte size and lipid content in order to reverse the inflammatory status of obese WAT [6–8]. Typical weight loss strategies consist of lifestyle modifications or, since 2017, glucagon-like peptide-1 (GLP-1) agonists. The former strategies are problematic as individuals have been shown to struggle following the exercise and/or diet interventions that have been prescribed, resulting in the “yo-yo” effect of weight loss/regain [9]. GLP-1 agonists are a revolution in weight management therapies; however, about ∼5% of patients taking GLP-1 agonists must be taken off due to severe side effects [10]. Most patients stop taking GLP-1 agonists after 2 years of therapy because they achieve sufficient weight loss or their diabetic symptoms resolve. Unfortunately, following cessation of GLP-1 therapy, it has been reported that patients will regain most of their weight after one year [11]. Therefore, the investigation of alternate weight management and weight-loss strategies needs to be continued. An alternate strategy for weight loss is through the induction of non-shivering thermogenesis (NST). NST represents the generation of heat from thermogenic adipose tissue to protect an individual’s core body temperature [12]. Activation of thermogenesis leads to increased energy expenditure through the catabolism of lipids and carbohydrates. This increased energy expenditure can decrease overall adiposity and, therefore, reduce systemic inflammation [13–17]. Currently, there is no FDA-approved therapy for weight management that mimics thermogenesis, yet our investigation provides a rationale that thermogenesis has therapeutic potential, especially for male individuals.

Baseline differences in thermogenic adipose have been noted in adult patients, showing that women tend to have more BAT than men do [18]. Recent correlative data from mice demonstrates sex-specific differences in energy expenditure, adipocyte mitochondrial function, and BAT activation [19]. While BAT represents the dominant thermogenic adipose, and it has been shown countless times that inguinal WAT (i.e., subcutaneous WAT) induces UCP1 during cold exposure or pharmacologic means. A significant controversy in the field persists as to whether thermogenic adipocytes are present and/or functional in perigonadal WAT (pgWAT), the visceral white adipose depot of mice. To this point, different studies have been reported conflicting data on whether UCP1 can be expressed in pgWAT [20–25].

Adipocytes are the primary cell type of adipose tissue; of which there are two primary types of adipocytes (white and brown). The primary role of white adipocytes is to store energy in the form of lipids; while the primary role of brown adipocytes is to metabolize lipids in order to generate heat to maintain a core body temperature [26, 27]. Yet, white adipocytes can also participate in thermogenesis when the animal is exposed to temperatures below their thermoneutral zone (TNZ) [21, 28]. The TNZ is defined as the temperature range when no additional energy expenditure is required to maintain core temperature. Adipose thermogenesis is activated by the sympathetic nervous system release of β-adrenergic receptor agonists [29]. Upon activation, these receptors increase intracellular lipolysis; resulting in the release of FFAs that can be taken up by mitochondria to fuel heat production [29, 30]. This heat production also requires coordination of transcription factor activation, including PPAR signaling, CIDEA, PRDM16, and PGC1α [31, 32]. These transcription factors bind to the promotors of mitochondrial transcriptional pathways and UCP1 to increase expression. UCP1 is located on the inner mitochondrial membrane and functions to allow protons to move down the electrochemical gradient generated by the electron transport system. This uncouples mitochondrial oxidative metabolism from the generation of ATP, resulting in increased heat production and active thermogenic adipocytes [26, 28, 32]. Interestingly, thermogenic adipocytes are less active as an individual’s BMI increases and tend to take on a white-like phenotype where they favor the storage of lipids [33]. Therefore, it can be concluded that BAT and active thermogenic adipocytes have the potential to be anti-obesogenic.

A valuable tool used by researchers and clinicians to analyze adipose depots in a non-invasive manner is CT imaging, which utilizes the attenuation of X-rays to reflect the density of tissues and generate cross-sectional images of the subject being imaged. The current gold standard to detect thermogenic adipose is the use of positron emission tomography-computed tomography (PET-CT) imaging. However, it requires the administration of radiolabeled glucose to detect active thermogenic adipose. Alternatively, the advancement of microCT (µCT) allows for access to CT technology on a smaller scale with increased achievable resolution. While contrast agents are commonly used to enhance x-ray attenuation and improve the identification and clarity of final µCT images, to our knowledge, a longitudinal method to monitor changes in adipose tissue based solely on variations in Hounsfield units— without the use of contrast agents—has yet to be developed. This method is a high priority for increasing efficiency in imaging and reducing bias in the identification of adipose tissues.

In this study, we developed a replicable method of detecting the adipose tissue changes and the formation of active brown adipose via non-contrast µCT with a streamlined segmentation strategy for the quantification over time. Additionally, we demonstrate that thermogenic adipocytes readily form in the BAT of obese mice but are formed to a lesser extent in the pgWAT of male mice once they undergo cold exposure. Surprisingly, we found that TN DIO female mice maintain a significant amount of thermogenic adipocytes in their brown adipose. In contrast to the male mice, the cold-exposed female mice were also capable of activating thermogenic adipocytes in their pgWAT despite BAT not showing additional gains in thermogenic adipocytes. Importantly, we provide evidence that cold exposure results in an alteration of adipose-secreted pro-inflammatory cytokines in both the WAT and BAT of male mice. Considering the potential for secretion of lipid metabolites, we observed no change in FFAs in the serum of the male or female mice between TN conditions and TS conditions. While the TS female mice showed increased glucose levels in their serum, we surprisingly detected a loss of glucose for the TS male mice. In conclusion, we have defined a way to differentiate and measure thermogenic adipose formation *in vivo* and describe a model of thermogenic induction that results in unique sex-based responses to cold exposure in obese mice.

## Results

### Whole body measurements of mice at varying environmental conditions

To determine how cold exposure affected thermogenic activation in obese adipose, mice were placed on a WD and housed at their TNZ of 30°C. The body weight of both male and female mice on WD was measured once per week (Fig. 1A). Male mice gained weight at a constant rate; upon reaching 35g, the mice were considered to be obese. TN male mice were then subjected to cold exposure via a thermalshift (TS) to induce thermogenic activation of the adipose. While a TS to either RT or 18°C resulted in a significant loss of body weight, the TS to 18°C cohort demonstrated the most significant decrease (Fig. 1A & 1C). However, these results were not recapitulated in the female cohort. The female cohort gained weight at a variable pace upon WD feeding at TN housing (Fig. 1B); even though they were subjected to the exact same conditions as the male mice. Moreover, in terms of body weight, the female mice did not show significant weight changes when exposed to either RT or 18°C (Fig. 1D); creating a sex-specific response between males and females.

**Figure 1.**
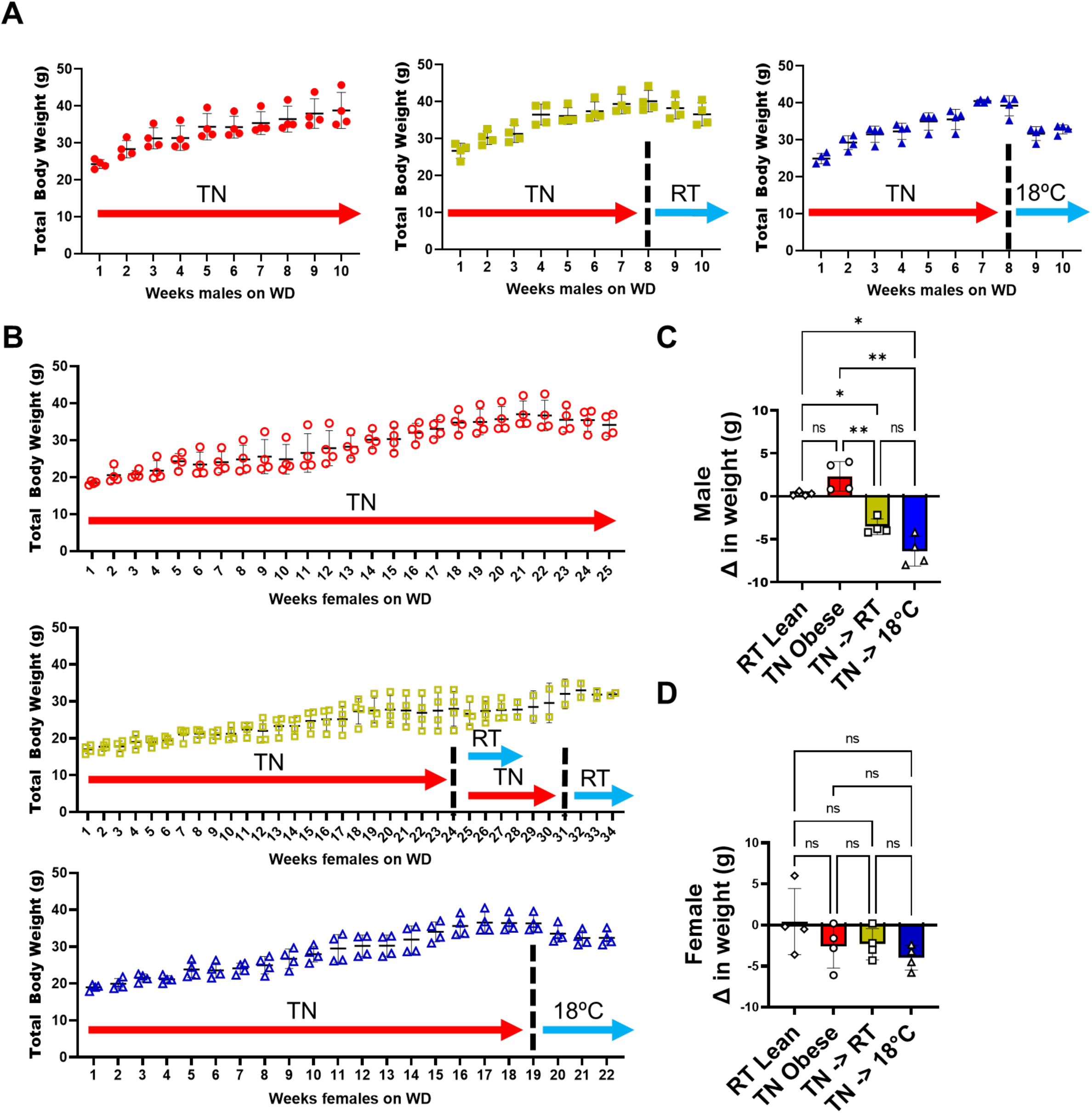
Weight loss in response to cold exposure in obese mice. Whole-body weight was measured once per week for **(A)** male and **(B)** female mice fed a WD. The change in weight before and after cold exposure in **(C)** male and **(D)** female mice fed a WD. Black dotted line denotes timepoint for the thermalshift, TS. *, P < 0.01; **, P < 0.001, ***, P < 0.0001, NS, P > 0.05; Brown-Forsythe and Welch one-way ANOVA of 4 biological replicates.

### A non-contrast µCT method of thermogenic adipose detection

Mice began serial µCT imaging once they reached obese status (except for room temperature lean mice), and then every seven days thereafter (Fig. 2). Initial imaging (Day 0) provided a baseline reading prior to changes in housing conditions for the relevant groups. All mice were imaged with the same parameters: one 4-minute scan at 36 mm field of view (FOV) at 90kV and 88uA with an Al 0.5 mm filter to verify brown fat depositions and confirm that signal present was not due to mis-registration of signal intensity due to motion artifacts, and a 2-minute stitched scan (for a total of 4-minutes) at 72 mm FOV to quantify whole body composition. Where possible, retrospective-respiratory gated scans were used; however, in select instances when gating was not successfully employed due to variable respiration rate during imaging, the averaged image was utilized. Images were windowed to facilitate differentiation between brown and white adipose tissue depositions with the window-level preset “CT-Abdomen” in 3D Slicer and modified further in rare instances by lowering the center of the windowing (Fig 2A).

**Figure 2.**
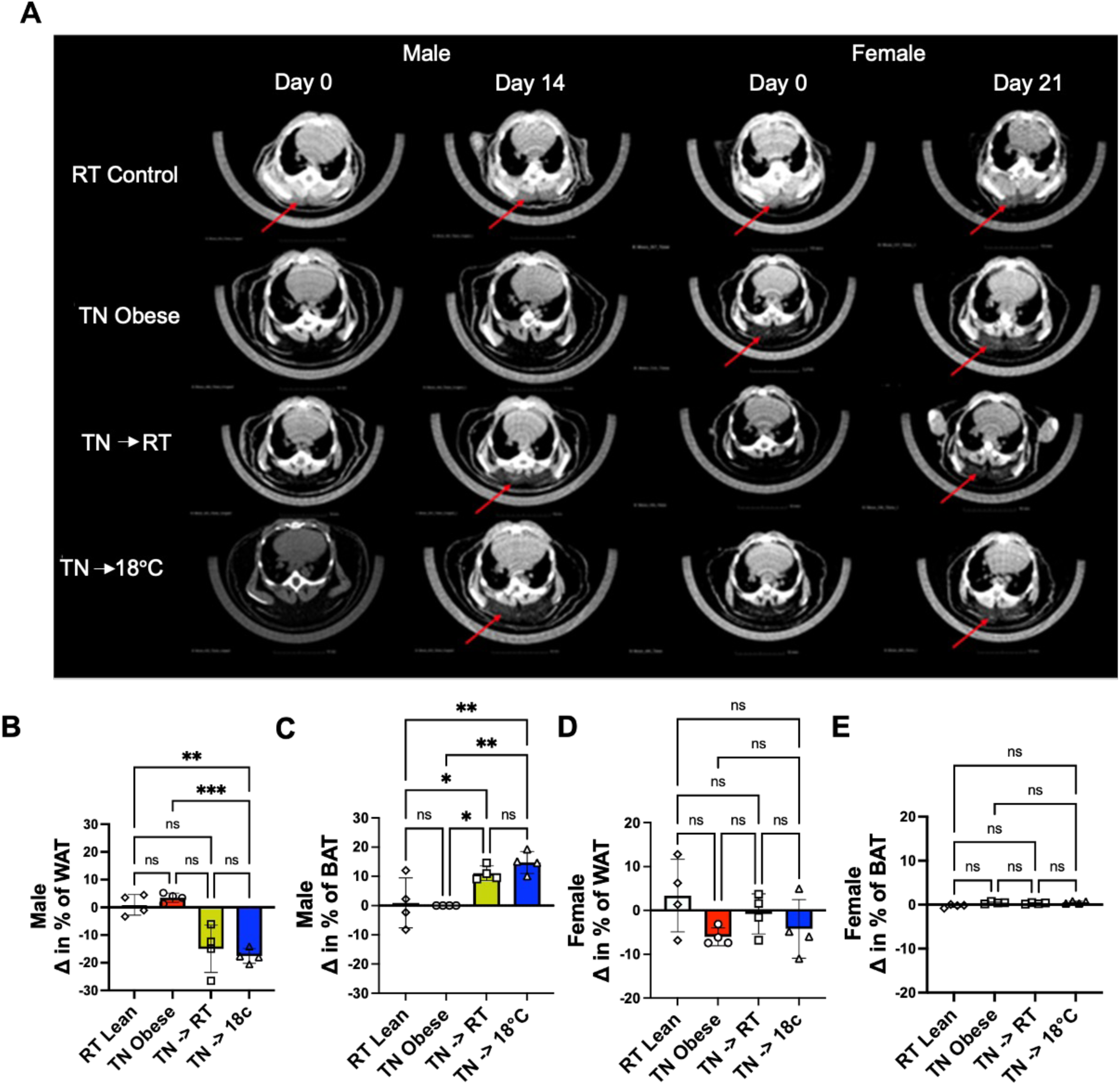
µCT imaging analysis in response to varying environmental temperatures. **(A)** Cross-sectional slices taken from each data series from the first (Day 0) and final imaging day (Day 14 for male, Day 21 for female) for each sex and environmental conditions (RT, TN, TN to RT, or TN to 18°C). Quantifications of the changes in WAT and BAT volume of male wild-type mice fed a WD before and after cold exposure **(B & C)**. Quantifications of the changes in WAT and BAT volume of female wild-type mice fed a WD before and after cold exposure **(D & E)**. *, P < 0.01; **, P < 0.001, ***, P < 0.0001, NS, P > 0.05; Brown-Forsythe and Welch one-way ANOVA of 4 biological replicates.

Different regions of anatomy were then segmented with arbitrary color assignments, though presets in 3D Slicer were used when possible, so long as they provided easy delineation between tissues. These segmentation values were then used to extract the scalar properties of the images such as CT value and volume for quantification and reference against histological data. Two imaging timepoints are represented for Day 0 versus Day 14 in male or Day 21 in female for each of the conditions (Fig. 2A). As seen in Figure 2A, male mice followed similar trends where TN obese starting points resulted in little to no brown fat depositions compared to RT control mice on initial imaging prior to any changes in housing conditions (rows: TN Obese, TN-> RT, TN-> 18°C and the first two columns on the left). Room temperature control mice (which stayed at RT throughout the study showed the highest HU values for brown fat measurements (average of 28 HU across all imaging days) with the densest appearance compared to other brown fat depositions across male mouse groups consistently across each timepoint with brown fat holding 18% of total body composition (Fig. 2A). Brown fat depositions for male mice moved to 18°C housing conditions showed alterations over time, with a peak at the mid-point followed by a minor decline at the endpoint, presumably as the rodents adapted to the new housing conditions (final average of 15% of total body composition) (Fig. 2C). HU values for rising brown fat depositions in male mice moved to 18°C were generally lower than those found in control mice indicating less density in this tissue compared to the room temperature BAT depositions, but did increase in CT value over time (average at day three of −70 HU) demonstrating the change in tissue characteristics due to tissue browning (Fig. 2A). Similarly, male mice that transitioned from TN to RT also saw a fat browning over time due to “room temperature” being a subthermal temperature for mice. Interestingly, the BAT depositions in male mice moved from thermoneutral housing to room temperature housing had increased density of brown adipose tissue compared to those moved to 18°C with an average CT value of approximately −60 HU (Fig. 2A).

Female mice varied compared to their male counterparts of the same grouping as seen in the two-columns on the right of Figure 2A. RT mice had similar results to the male mice with consistent, dense depositions of brown fat across each imaging time point with the highest HU values averaging at 30 HU across all timepoints (Fig. 2A). Variances between brown fat depositions appeared for some mice in the TN cohort, with visualization and possible segmentations of these brown fat depositions on the first imaging day in some individuals – however it is worth noting, that female mice also had more difficulty reaching obesity (Fig. 1B). Generally, these regions which are visualized as depositions of BAT in TN female mice were diffusively distributed across voxels in these regions and were largely removed during the smoothing process of applying opening and closing kernels to solidify the different segmentations. Alternative segmentation strategies were employed to assess the HU values for these regions, and they reflected the lower end of reasonable CT values for brown fat (−70 HU) given average HU values for brown fat across all groups. Interestingly, at endpoint, TN DIO female mice, on average, had denser allocation of BAT with increased HU values over time (−35 HU at end point) when measured with µCT (Fig. 2A). Female mice which changed housing conditions over time, such as TN to RT and TN to 18°C both saw BAT depositions at endpoint, yet similarly to the TN DIO female mice, most also contained brown adipose at the starting point of imaging with little differences in HU value and % composition in BAT over time; although female mice moved to 18°C saw slightly elevated HU values (average −29 HU at end point) compared to those shifted to RT (average −63 HU) (Fig. 2A). These µCT findings in female mice indicate more substantial dense blocks of different adipose tissue types, while clear boundaries are seen in their male counterparts.

### µCT based quantification of thermogenic adipose

To assess the extent of thermogenic activation over time in mice, we used our µCT method of analysis outlined in Figure 2 to quantify total body WAT and total BAT (Fig. 2B-E). We observed a significant decrease in total WAT in the male mice exposed to RT or 18°C when compared to the DIO mice that stayed at TN (Fig. 2B). In contrast, the WAT in the female mice did not significantly change between the obese mice that stayed at TN and the female mice exposed to either RT or 18°C (Fig. 2D). Quantification of BAT also showed a similar dichotomy between the two sexes. Specifically, the cold-exposed male cohort showed a significant increase in brown adipose content (Fig. 2C). In comparison, the female mice did not show any significant changes in their BAT content from imaging analysis (Fig. 2E). In the assessment of this data, the male mice adipose depots showed significant changes during cold-exposure whereas female mice fail to display detectable changes in body mass or µCT adipose composition analysis.

### Histological analysis of adipose tissues

To validate that tissue-specific changes in the adipose were consistent with the µCT imaging analysis, we histologically assessed the pgWAT and interscapular BAT (iBAT) depots for both sexes. Hematoxylin and eosin (H&E) staining revealed an increase in the presence of white-like adipocytes in TN iBAT (Fig. 3A, top images) compared to the classically activated BAT phenotype in mice fed a CD and housed at RT. Upon cold exposure from TN to either RT or 18°C, male mice demonstrate reversion to a more classically activated BAT signature in their iBAT; this resembles the lean RT brown adipose. When assessing the pgWAT of DIO males (Fig 3A, bottom images) at TN compared to mice fed CD and housed at RT, there was a hypertrophy of the white adipocytes. Upon cold exposure from TN to either RT or 18°C, male mice demonstrated a modest reduction in adipocyte size, yet mostly retain phenotypically large adipocyte sizes in their pgWAT. Because the µCT demonstrated significant percentage changes for total WAT and total BAT between the TN DIO adipose and the TS to 18°C, we quantified the area of adipocyte size in the pgWAT of these two environmental temperature exposures (Fig. 3B), which revealed a trending but non-significant decrease in adipocyte size in the male mice that underwent TS to 18°C versus TN DIO male mice.

**Figure 3.**
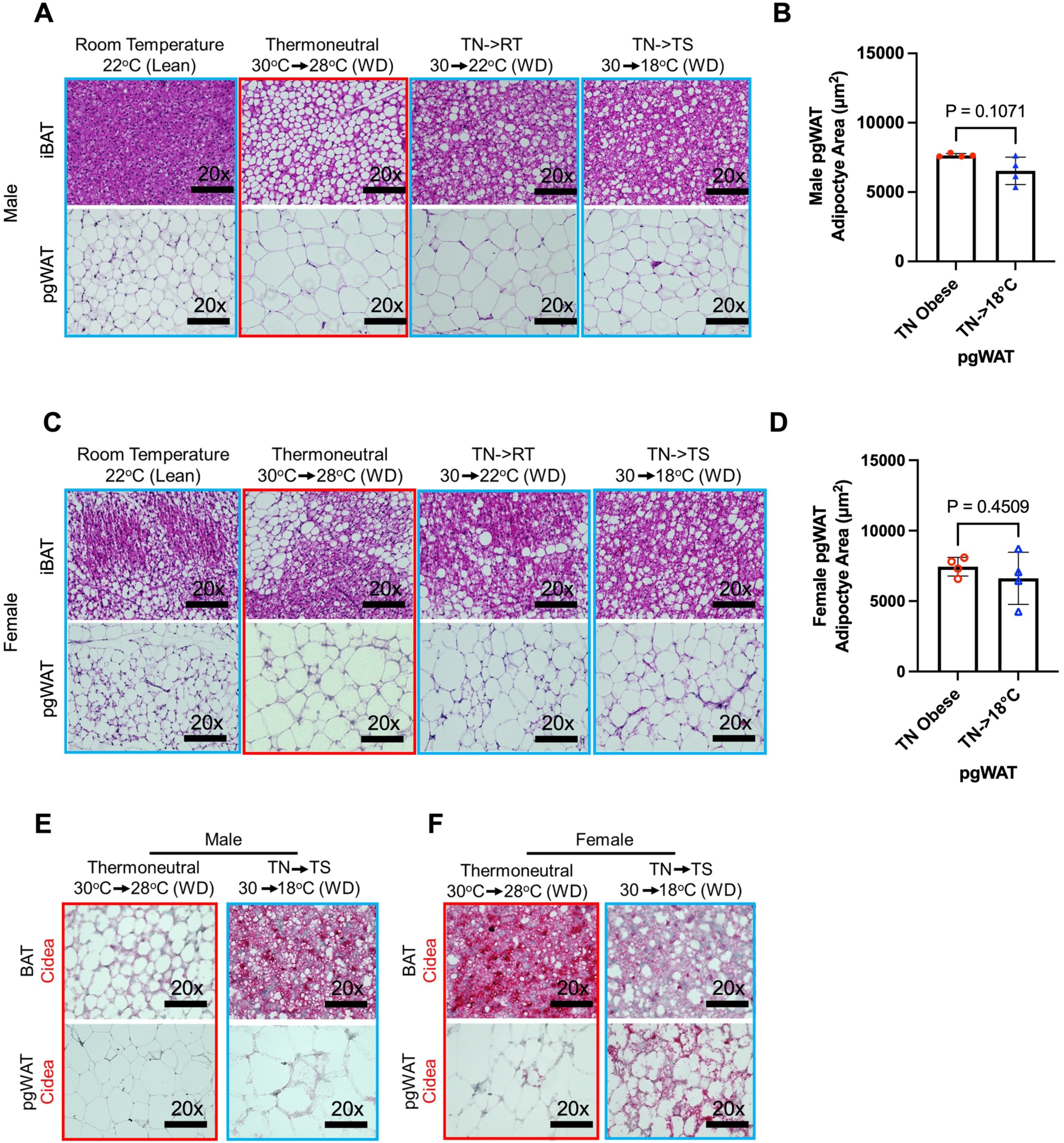
Histological analysis of sex-dependent changes in thermogenic adipose formation. Hematoxylin and eosin stain of WAT and BAT at different temperature exposures of **(A)** male and **(C)** female mice (representative images, n = 4). Adipocyte area (µm^2^) was quantified in the pgWAT of TN DIO and TS to 18°C of both **(B)** male and **(D)** female mice (4 images per biological replicate were measured). RNAscope was used to probe for *Cidea* transcripts in **(E)** males and **(F)** females of their WAT and BAT of TN or TS to 18°C cohorts (representative images, n = 4). NS, P > 0.5. Welch’s t-test.

Next, we assessed the female mice, which showed phenotypically heterogeneous iBAT in all cohorts of mice and no noticeable changes in composition across groups. Intriguingly, the adipose tissues in the female mice do show more heterogeneity in cellular histology than the male mice. There was a mixture of white-like as well as classic thermogenic adipocyte signatures that were located in the iBAT of all cohorts of female mice (Fig. 3C). While pgWAT adipocyte size increased from RT lean females compared to TN DIO female mice, there was little to no phenotypic change in their white adipocyte size when comparing TN female mice to the female mice that underwent TS to RT or to 18°C (Fig. 3C & 3D). Coincidently, H&E staining revealed that there were thermogenic-like adipocytes in the pgWAT interspersed among the classic white adipocytes present in all cohorts of female mice (Fig. 3C). These histological assessments revealed phenotypic differences in temperature as well as highlighted differences between the two sexes in their respective adipose tissues.

To understand the molecular alterations associated with adipose samples, we assessed markers indicative of thermogenic adipose via RNAscope and immunohistochemistry. We found that the thermogenic marker *Cidea* was had low expression in TN DIO male brown adipose, yet *Cidea* was expressed higher in the BAT of the male mice that underwent TS to 18°C (Fig. 3E). Additionally, another thermogenic marker UCP1 (data not shown) was expressed in a similar manner as *Cidea* for the male mice. Interestingly, the female mice were positive for the presence of *Cidea* in BAT of TN DIO mice cohort and exhibited an increased prevalence of *Cidea* in the pgWAT of female mice TSed to 18°C (Fig. 3F), which was not observed in male pgWAT at either TN or 18°C (Fig. 3E,F). As expected from the imaging analysis, male mice showed thermogenic adipose in their iBAT; while the female mice preferentially increased the thermogenic adipose in their pgWAT rather than in their iBAT. Importantly, the pgWAT thermogenic activation was not observed with our imaging analysis.

### Free fatty acid and glucose levels in cold exposed mice

To understand if systemic bioenergetics were changed in correlation with adipose alterations, serum was collected from mice at the study endpoint to measure FFAs and glucose levels in both female and male mice. Cold exposure of 18°C had no significant effect compared to TN housing on serum FFAs, but there was a notable difference between the sexes (Fig. 4A). Surprisingly, the serum glucose was decreased in the male mice exposed to 18°C, while it was increased in the female mice exposed to 18°C (Fig. 4B). Together, this data demonstrates that cold exposure has a unique sex-dependent glucose response between thermalshifted mice.

**Figure 4.**
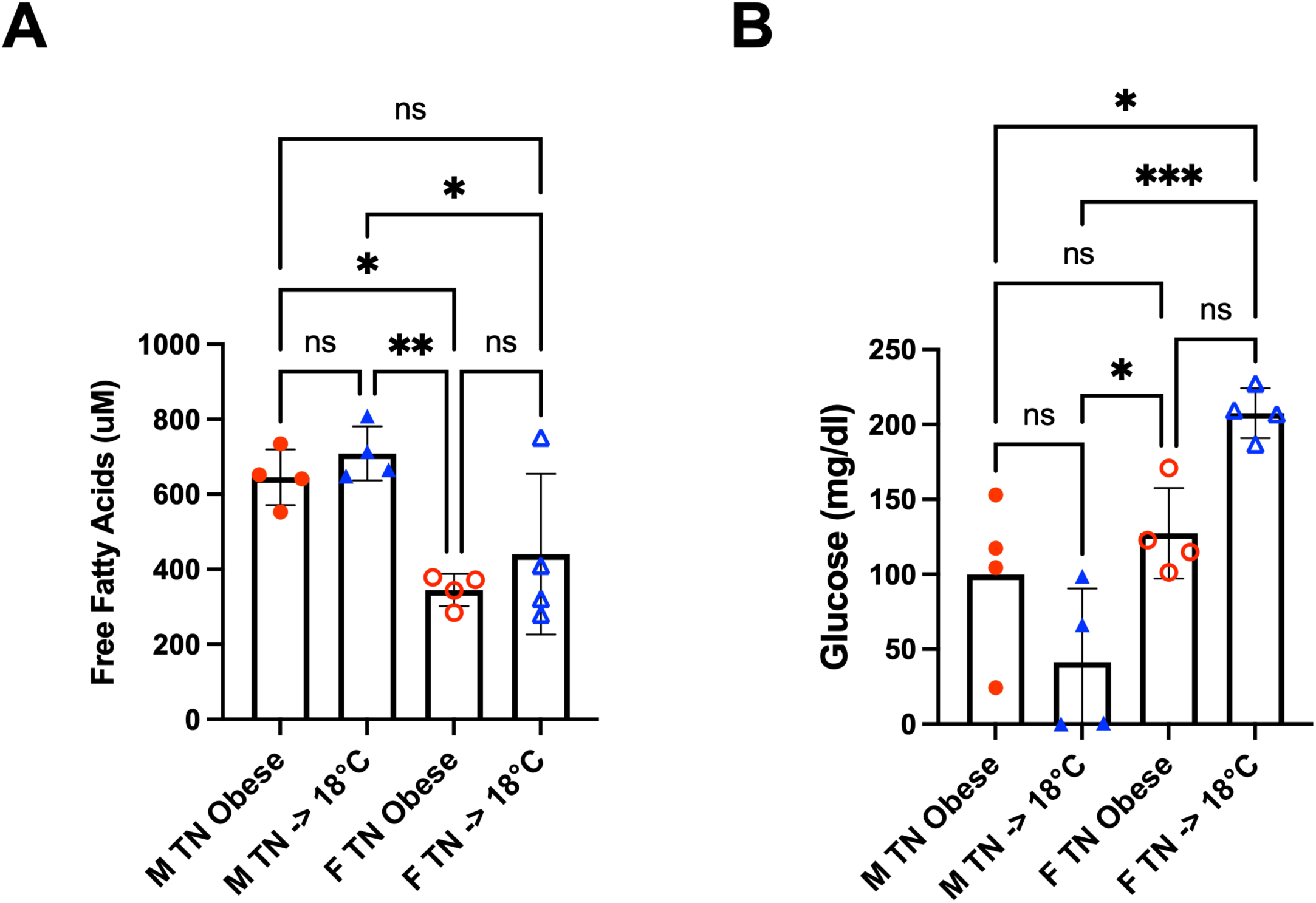
Analysis of serum metabolites in the context of diet and environment. Serum free fatty acid (µM) levels **(A)** and serum glucose (mg/dl) levels **(B)** in male (M) and female (F) mice fed a WD that stayed at TN or were cold exposed to 18°C (TN->18°C). *, P < 0.01; ***, P < 0.001, ****, P < 0.0001, NS, P > 0.05. Brown-Forsythe and Welch one-way ANOVA of 2 technical replicates per biological replicate of 4.

### Cold exposure affects cytokine secretion from adipose

Obese individuals typically display low-grade chronic inflammation of the adipose. Therefore, to determine if a TS to 18°C is enough to change the pathological inflammation, we performed label-free proteomic mass spectrometry based analysis of the conditioned media (CM) from adipose tissue explants of these mice (Fig. 5). When comparing TN DIO male mice adipose tissues, we found a decrease in secretory cytokine ligands including CCL5, CXCL10, CCL2, and CXCL1 for the WAT of 18°C male mice when compared to obese WAT in TN male mice (Fig. 5A); yet we found a loss of CXCL3, IL-6, and CXCL5 for the BAT of 18°C male mice compared to obese BAT in TN male mice (Fig. 5B). These cytokine changes in the WAT and BAT suggest that cold exposure reduces the overall pro-inflammatory response.

**Figure 5.**
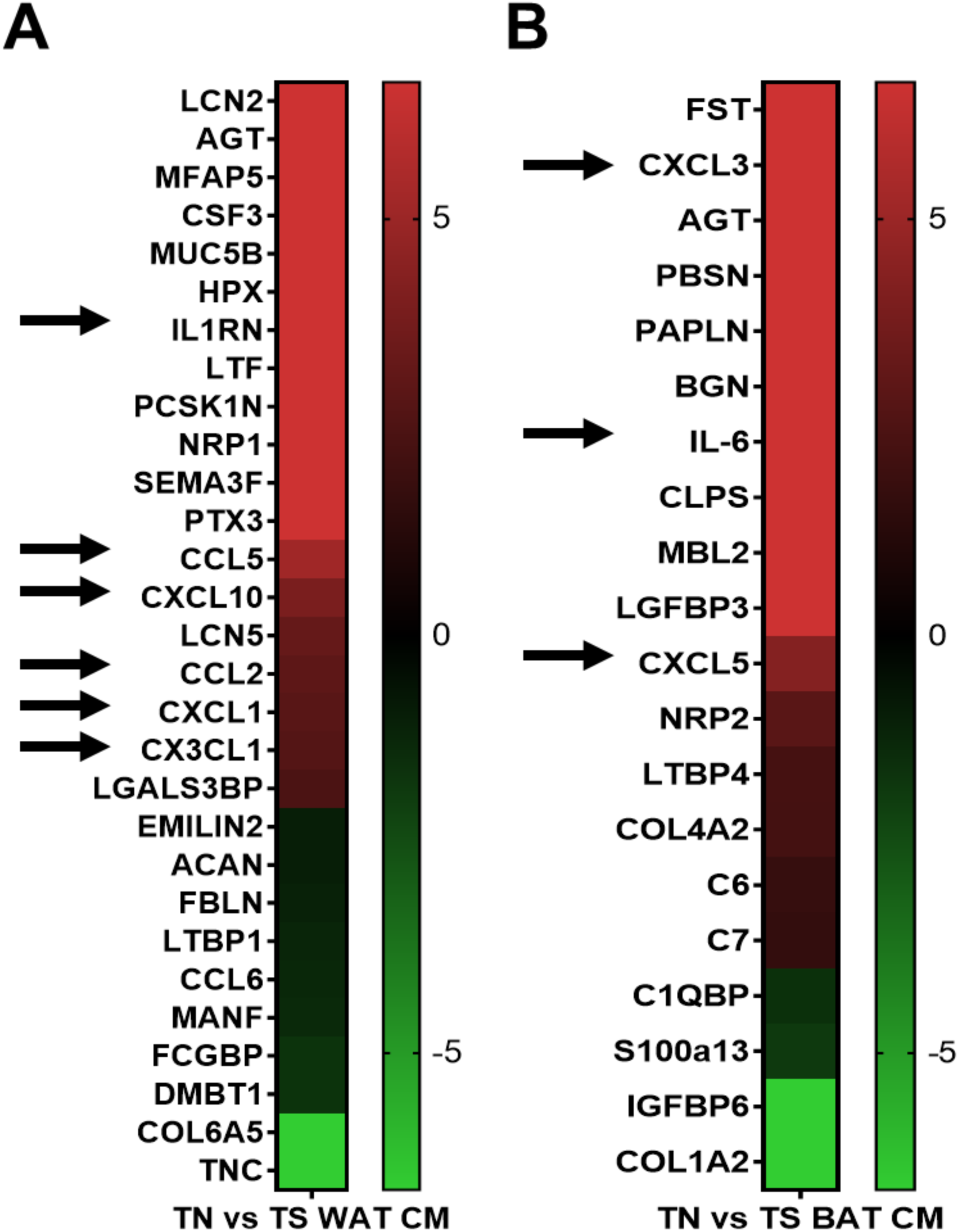
Identification of inflammatory mediators in adipocyte conditioned media (ACM) between TN and TS conditions. Using quantitative DDA proteomics, secreted factors were probed in DIO obese male pgWAT CM and pgWAT TS to 18°C CM **(A)**. The abundance ratio and log2(AR) were calculated for each protein of each comparison in order to determine the change in protein abundance; it was considered significant if P < 0.03. Using quantitative DDA proteomics, secreted factors were probed in DIO obese male iBAT CM and iBAT TS to 18°C CM **(B).** The abundance ratio and log2(AR) were calculated for each protein of each comparison in order to determine the change in protein abundance; it was considered significant if P < 0.03. Black arrows denote known pro-inflammatory cytokines. Representative of 3 technical replicates for each biological replicate (n = 4) per group.

## Discussion

The physiological state of adipose tissue is dynamic, responding to both nutrient levels in the body and changes in environmental temperatures. Adipose tissue assesses the bioavailability of nutrients in conjunction with energy balance and energy expenditure (EE) to determine whether to store, release, or burn energy-rich lipids. Understanding the intricate relationship between lean and obese states is essential, particularly in the context of the current obesity epidemic. A key aspect of studying adipose tissue involves monitoring the physiological shifts in adipocytes between white and brown subtypes. These adipose subtypes can be influenced by dietary changes or environmental conditions. Chronic consumption of excess nutrients tends to induce obesity, driving adipocytes toward a white-like phenotype. In contrast, exposure to cold temperatures or dietary restriction can encourage a brown-like phenotype in adipocytes, due to its higher metabolic rate. Therefore, promoting a transition of adipocytes to a more metabolically active phenotype and enabling them to utilize excess nutrients for energy rather than simply storing them, may provide novel methods to reduce the burden of obesity.

The concept of thermogenic activation to alter obese pathological adipose into a healthier brown-like phenotype has previously been implicated. For instance, Van Der Stelt et al. used a DIO model where they switched DIO male mice from TN to RT for 5 days [34]. They performed bulk RNAseq on pgWAT, which demonstrated that *Ucp1* was undetectable after RT exposure and failed to demonstrate any other significant changes in other thermogenic markers. Interestingly, however, they found that glucose in DIO male mice was down when switched to RT and that the FFAs were not changed between the two temperatures; but the TGs were down in the cold exposed cohort. In comparison, our data indicates that a duration of 14 days of chronic cold exposure is required for the complete development of thermogenic BAT. Additionally, we found that UCP1 induction in the pgWAT of male mice occurs at a significantly lower temperature of 18°C, providing further evidence of the presence of UCP1 in pgWAT. This was further corroborated by the female mice showing extensive UCP1 induction in their pgWAT even though all cold-exposed mice remained on WD throughout the study. Additionally, Zheng et al. displayed UCP1 induction in pgWAT when lean male mice were given a *β*_3_-adrenergic receptor agonist, CL316,243, for seven days [79]. The observation that females exhibited progressively greater amounts of thermogenic adipose tissue in their pgWAT is surprising, and the mechanism regulating this phenomenon remains unclear. In the context of energy expenditure (EE), Morris et al. recently demonstrated that EE is higher in female mice on a WD due to differences in fat and fat-free mass [35]. In our studies, female mice took a drastically longer time to become obese than males did, which further adds evidence to females maintaining higher thermogenic adipose and feasibly higher EE. It is important to consider the broader context of how male and female mice respond differently to a WD; current data supports a sex-specific difference in adipocyte responses.

Pro-inflammatory responses are typically associated with the development of obese adipose. Therefore, changes in adipose inflammatory factors between TN and TS were investigated (Fig. 5). Our analysis demonstrated a loss in CCL5, CXCL10, CCL2, and CXCL1 in the WAT and loss of CXCL3, IL-6, and CXCL5 in the BAT after the subsequent thermoshift. CCLs, CXCLs, and IL-6. With the rewiring of the inflammatory signature in the TS cohort, we see concurrent shifting of cytokines reflect changes from pro-inflammatory immune cells to anti-inflammatory immune subsets. Interestingly, Van Der Stelt et al. in their bulk RNAseq analysis of DIO male mice switched to RT for five days found that M1 macrophages were decreased in the pgWAT, confirming our results and demonstrating an inflammatory shift occurs in WAT during cold exposure [34]. Therefore, the systemic inflammatory signature of obesity can be reduced by cold exposure through formation of thermogenic adipose as shown by us and others.

Metabolic dysfunction is a hallmark of obesity and can manifest into debilitating illnesses; cold exposure could have an impact on renormalizing metabolism in the setting of obesity. Hao et al. showed glycolytic genes were up in male BAT during exposure to 4°C for 8 days, suggesting that BAT heavily relies on glucose uptake to maintain heat production as well as that it can be used for fatty acid synthesis [36]. Additionally, Iwen et al. demonstrated that glucose was depleted in the plasma of 15 healthy men after 100 minutes of cold exposure to 18°C [37]. Our results demonstrated that serum glucose was depleted in DIO male mice exposed to 18°C but increased in DIO females exposed to 18°C (Fig. 4B). This may be indicative of impaired glycolysis in BAT of the female mice and that the increase in thermogenic adipocytes in WAT of the female mice does not compensate for this impairment. The other fascinating finding was that there was a lack of shift for serum FFAs between the two temperatures for either sex; however, there was a distinct basal difference between male and female mouse cohorts (Fig. 4A). This could suggest that male mice have a propensity to use more FFAs than female mice, possibly due to having more muscle mass overall. Alternatively, this could mean that males consumed more of the WD on average when compared to females owing to their increased rate of weight gain. Unfortunately, we did not record food consumption, but Morris et al. demonstrated that TN female mice consume less WD over a span of 7 days when compared to TN male mice consuming the same WD [35]. In regard to FFAs, our results don’t agree with what Iwen et al. also discovered, which was that specific plasma fatty acids are altered in the fifteen men cold exposed; but this was in healthy men and it is unknown if the FAs would be altered if they had a BMI above normal weight let alone what happens in women [37]. Moreover, Khedoe et al. show that cold-exposed BAT burns intracellular TGs and needs to take up lipoproteins and FFAs derived from lipolysis after intracellular TGs are burned [38]. This could also mean that our mice cohorts had enough lipids to burn intracellularly and did not need exogenous FFAs.

The non-contrast µCT parameters used in this study are available on most preclinical scanners. They can effectively detect and analyze thermogenic activation in the adipose tissues of both male and female obese mice. This approach builds on previous work by reducing user bias and decreasing analysis time through a segmentation strategy based on HU values, rather than relying on manual segmentation [39]. This analysis revealed a sex-specific thermogenic response between males and females, which is a novel discovery that may lead to insights into developing personalized therapies. To that end, cold exposure remains an exciting area of investigation for manipulating adipose in a non-invasive manner that has little to no side effects, as seen with pharmacological methods.

Limitations of the study: Although out of the scope of this study, important future investigations should focus on how the liver responds to cold exposure with regard to metabolite trafficking. It is unclear if the liver develops steatosis during cold exposure or if it contributes to FFA release, which could explain why serum FFAs did not change. It is also unknown whether increased gluconeogenesis in the liver is the reason females have higher serum glucose during cold exposure, as muscle can be impacted by cold exposure, but it was out of the scope of this study and, therefore, still needs to be analyzed in this context. Importantly, an in-depth analysis of the usage of amino acids as a fuel source for thermogenic adipose during cold exposure needs to be performed. Kovanicova et al 2021 shows in humans exposed to acute cold of 15mins in ice water had no change in circulating glucose levels but the alanine, aspartate and glutamate metabolic pathways were significantly affected [40]. Also, Okamatsu-ogura et al. found that glutamine is utilized in BAT in a UCP1-dependent manner during acute cold exposure [41]. To that end, we hypothesize that certain amino acids do have a role in WAT and BAT during chronic cold exposure, but it was not formally tested in our study. Lastly, energy expenditure, food intake, and menstrual cycles were not measured; these could provide alternate explanations for the differing effects of thermogenic adipose formation between male and female mice.

Imaging studies faced certain limitations – such as variance in image noise that was dependent upon sample size. While all subjects were imaged with comparable parameters, the smaller size of female mice pre-disposes these datasets to a decreased signal-to-noise ratio. While this experiment did not delve into complex post-processing techniques, future iterations could include post-processing steps to decrease the incidence in noise in the final image and increase image quality. Additionally, due to variances in size, smoothing kernels used throughout the various segmentations had larger impact on female mice due to the decreased total number of populated pixels in the cross-sectional slice. This led to slight under-estimations for final segmentations in females compared to males as each smoothing kernel covered a larger area respective to the total area occupied by the sample. Additionally, CT values are known to fluctuate with respect to hydration, and this fluctuation was not accounted for throughout imaging [42, 43]. However, it is important to note that mice were never separated from a source of water throughout imaging, and imaging times were less than 10 minutes per mouse with low-likelihood of induced dehydration. Due to utilizing a non-contrast form of imaging, muscle and organs presented at close HU values that were not accurately separable through our given segmentation strategy. For consistency these components were grouped under the heading “muscle” in the percent body composition but present an additional complication in analysis. Finally, all images were segmented through a threshold-technique, with rare instances of flood-filling used to delineate different tissues more rapidly. This was done remove user bias and instead focus primary on measured CT values in assigning voxels and calculating body-percentages. Smoothing kernels were then used to remove small assignments that were assumed to be due to noise or motion artifacts. However, the utilization of these kernels makes it possible to incorrectly classify a given pixel by removing its initial designation. To avoid this, a secondary segmentation strategy was employed to exclusively utilize thresholding to segment all components of anatomy and compare findings – indicating comparable values across groups with minor variances.

## Materials and methods

### Animal Studies

All animal studies were performed in accordance with the approved IACUC protocols at the University of Kansas Medical Center.

### Diet-induced Obesity (DIO)

6-8 week old C57BL/6J male or female mice were obtained from Jackson Research Laboratories, separated into appropriate cages and fed *ad lib* either a low-fat diet (5001, LabDiet: 13.5% calories from fat, 58% from carbohydrates, and 28.5% from protein) or Western-Style Diet (WD) [Teklad-Envigo InotivCo TD.88137] (adjusted Calories Diet (42% kcal from fat, 34% sucrose by weight) until 30g for females or 35g for males. Body wight was monitored once per week. During the feeding study, mice were either housed at standard RT of 22°C or TN of 30°C [Alternative Design Manufacturing & Supply Solace Zone Thermal Zone Technology. For mice that were exposed to 18°C, Sable Systems International Promethion Core CAB-16 Environmental Control Cabinets were used where mice were housed for 2 to 4 weeks.

### µCT imaging

All mice were imaged in the Quantum GX2 microCT at 90 kV and 88 uA with an Al 0.5 mm filter with ring reduction on. Image data was stored in a 512 x 512 matrix with voxel sizes of 72 mm for the 36 mm field of view (FOV) scan and 144 m for the 72 mm FOV. Prior to imaging, mice were removed from caging and placed into an anesthesia induction chamber at 4-5% isoflurane for knockdown. Once sufficiently anesthetized, mice were placed supine on the imaging cradle with their nose in a nosecone, upper limbs restrained alongside their head for a clear view of their diaphragm for gating, and their hind limbs gently restrained along the cradle to limit motion artifacts. The mice were centered in the gantry and imaged with two acquisition series. For the high-resolution scan with the 36mm FOV, the mice were centered with upper body in the imaging frame and a smaller field of view centralized over the diaphragm for retrospective respiratory gating. The secondary scan was taken at 72mm FOV, where the base of the skull and start of the tail positions were annotated as mm-along the cradle with these positions utilized to start and stop a 4-minute stitched scan for fully body data set. All centering of mice was performed at 90kV and 40 uA to minimize accumulated radiation, all radiation was limited to 2.5 Gy per imaging day.

After imaging, data is stored as DICOM files and then converted to NRRD for data consolidation and organization. High-resolution and standard-resolution data sets were analyzed separately, with full-body assessments being made from the whole-body series, and the high-resolution data set was predominantly utilized for confirmation of brown fat location behind the scapula. Images were segmented and analyzed in 3D slicer for brown and white fat quantification. The images were cropped to eliminate erroneous background and remove ringing artifacts from ear tag presence in the imaging where applicable. In the first and last slice of the image series, the cradle is segmented and then filled between slices. Bone was segmented through thresholding with average values of 1230-1300 HU. Segmentations were then selected to apply only to the outside of current segmentation values. Muscle was segmented through thresholding with average values of 90-100 HU, followed by an opening kernel of 5 x 5 or 3 x 3, depending upon the size of the sample. Following the opening kernel, a closing kernel of 3 x 3 was applied to close the segmentation. Lungs were segmented by initially level tracing the centermost region of the lungs in the coronal view to outline both lobes as well as the trachea. Flood filling with intensity tolerances of 170-200 and neighborhood sizes of 5-8 (depending upon the degree of adherence to the region of the lungs) was applied to segment the entire airway appropriately. The average intensity of the lungs was approximately (−320) – (−345) HU. Brown fat depositions were initially visualized behind the scapula of the mice and thresholded with average values of (−34) – (−64) HU depending upon sex and degree of browning (control mice having the highest HU values for brown fat). Similarly to the muscle segmentation, a series of 3 x 3 or 5 x 5 filters were used to open and close the segmentation. Finally, the white fat was segmented with average HU values of (−140) – (−145) HU. White fat segmentations were then closed with a 3 x 3 kernel. Some instances of segmentation utilized larger kernel sizes up to 7 x 7 pixels (dependent upon total image size, generally reserved for post-mortem imaging and large samples maximally occupying the field of view) for muscle and brown fat. After segmentation, the volume and HU value was extracted with 3D Slicer and collated for analysis. All average Hounsfield Units were determined by total measurements across all groups, specific values by group and housing condition varied according to tissue density within subtype.

### Histology

As described previously [44], tissues were fixed immediately after collection in 10% formalin at room temperature for 16-32hrs and transferred to 70% ethanol before paraffin processing, embedding, and sectioning. Slides were rehydrated and stained with either Hematoxylin and Eosin (Electron Microscopy Sciences, #26252-01 and #26252-02), processed for immunofluorescence (IF) staining, or for RNAscope™. For IF staining, tissue was sectioned at 8uM or 10uM for brown adipose tissue and for white adipose tissue, where they dried at 37°C overnight. Sections were permeabilized using TBS with 0.05% Triton X-100 (Sigma Aldrich, #T9284) for 15 minutes. Antigen retrieval was performed by microwaving slides in 10mM tris-EDTA solution (50mM Tris pH 8.8, 1mM EDTA, and 0.05% Tween 20). Tissue sections were blocked in donkey serum-containing blocking buffer (2M Tris pH 7.4, 1M MgCl, 0.05% Tween 20, 2% BSA, 5% Donkey Sera) for 1 hour at room temperature before adding primary antibodies in a 1:1 ratio of donkey serum-containing blocking buffer and TBS for overnight incubation at 4°C. Slides were then incubated with secondary antibody in blocking buffer containing (5µg/mL) Hoechst 33342 (Invitrogen #H3570) for 1 hour at room temperature. Slides were mounted using Invitrogen ProLong Diamond Antifade Mountant (Thermo Fisher #P36961) for IF slides and Permount mounting medium (Thermo Fisher #SP15-100) for H&E slides. All slides were imaged using an EVOS E5000 (Thermo Fisher #AMF5000) microscope. Primary antibody used for IF Staining was UCP1 (R&D research MAB6158) and secondary antibody used was Alexa-Fluor 488 donkey anti-mouse IgG (H+L) (#A21202). *Cidea* transcript was detected via RNAscope™ according to ACDbio’s protocol. In brief, we utilized the RNAscope™ 2.5 HD Detection Reagents-RED kit (Cat No. 322360) with the HybEZ™ II Hybridization System for Manual Assays which is comprised of HybEZ oven (PN 321710/321720), a humidity control tray (PN 310012), and HybEZ Humidifying Paper (PN 310025), EZ-Batch Wash Tray (PN 321717), and the EZ-Batch Slide Holder (PN 321716). Additionally, to successfully carry out the manufactures protocol we used the accessories ImmEdge® Hydrophobic Barrier Pen (Cat. No. 310018), and the EcoMountTM Mounting Medium (Cat. No. 320409). All tissues were fixed with 10% formalin at RT for 16-32hrs transferred to 70% ethanol before paraffin processing, embedding, and sectioning at 5uM per section. To detect *Cidea* transcript we used the RNAscope™ Probe-Mm-Cidea-C2 (Cat No. 455441-C2). For antigen retrieval, Black & Decker Steamer was used and set according to ACDbio’s protocol. For adipocyte area quantification, slides were imaged with EVOS M5000 (Invitrogen) and quantified using ImageJ.

### Adipose Conditioned Media

Adipose tissue is extracted, weighed, and the adipose collected will be washed for 2 hours in serum free media (SFM) containing anti-anti and 10ug/ml of ciprofloxacin. After two hours, the wash media will be gently decanted and fresh SFM containing anti-anti and 10ug/ml of ciprofloxacin will be added at a ratio of 125mg of adipose tissue per 1ml of SFM. The adipose will incubate in the SFM for 24hours, collected into a sterile tube, and the adipose conditioned media (ACM) will be spun at 5,000xG for 3 minutes. After centrifugation, the fat cake will be carefully removed, and avoiding the pelleted debris, the ACM will be sterile filtered with a 20uM filter and aliquoted to put into −80 storage until needed for downstream applications. When used for assays, a working solution of 1:3 to 1:5 dilution with preferred media will be prepared.

### Sample preparation for LC-MS/MS analysis

As described previously [45], adipose conditioned media was collected as described above and were reduced by adding 50mM TCEP to a final concentration of 5mM followed by incubation at 55°C for 30 minutes. Cysteines were alkylated by the addition of 375mM IAA to a final concentration of 10mM followed by incubation at room temperature in the dark for 30 minutes. Ice cold acetone was added at a 1:5 ratio followed by incubation at −20°C overnight. After precipitation, samples were centrifuged at 14000 x g at 4°C for 10 minutes to pellet the proteins. The supernatant was removed, and proteins were allowed to air dry on the bench top for 15 minutes. The proteins were resuspended in 100ul of 50mM TEAB pH 8 with 2mM CaCl2 and proteins were digested by adding 500ng of trypsin and incubating overnight at 37°C at 500 RPM (Thermomixer, Eppendorf). Digestion reactions were quenched by the addition of 10% formic acid to a final concentration of 1%. Digested samples were centrifuged at 10,000 x g for 10 minutes to remove particulates and supernatant was transferred to a fresh tube for LC-MS/MS analysis. Peptide concentrations were measured using a Nanodrop spectrophotometer (Thermo Scientific) at 205 nm.

### LC-MS/MS detection and data analysis

As described previously [45], samples were injected using the Vanquish Neo (Thermo) nano-UPLC onto a C18 trap column (0.3 mm x 5 mm, 5 µm C18) using pressure loading. Peptides were eluted onto the separation column (PepMap™ Neo, 75 µm x 150 mm, 2 µm C18 particle size, Thermo) before elution directly to the MS [45]. Briefly, peptides were loaded and washed for 5 minutes at a flow rate of 0.350 µL/min at 2% B (mobile phase A: 0.1% formic acid in water, mobile phase B: 80% ACN, 0.1% formic acid in water). Peptides were eluted over 100 minutes from 2-25% mobile phase B before ramping to 40% B in 20 min. The column was washed for 15 min at 100% B before re-equilibrating at 2% B for the next injection. The nano-LC was directly interfaced with the Orbitrap Ascend Tribrid MS (Thermo) using a silica emitter (20 µm i.d., 10 cm) equipped with a high field asymmetric ion mobility spectrometry (FAIMS) source. The data were collected by data-dependent acquisition with the intact peptide detected in the Orbitrap at 120,000 resolving power from 375-1500 m/z. Peptides from samples with a charge of +2-7 were selected for fragmentation by higher energy collision dissociation (HCD) at 28% NCE and were detected in the ion trap using a rapid scan rate. Dynamic exclusion was set to 60s after one instance. The mass list was shared between the FAIMS compensation voltages. FAIMS voltages were set at −45 (1.4 s), −60 (1 s), −75 (0.6 s) CV for a total duty cycle time of 3s. Source ionization was set at 1700 V with the ion transfer tube temperature of 305°C. Raw files were searched against the mouse protein database downloaded from Uniprot on 09-26-2023 using SEQUEST in Proteome Discoverer 3.0 [46]. Abundances, abundance ratios, and p-values were exported to Microsoft Excel for further analysis.

### Statistical analysis

The change in total body weight was calculated by weight at the beginning and endpoint of µCT imaging with 4 biological replicates for each experimental condition and sex, statistical significance was calculated with Brown-Forsythe and Welch one-way ANOVA (*, P < 0.01; **, P < 0.001, ***, P < 0.0001, NS, P > 0.05). Changes in adipose were quantified by µCT as described above; statistical significance was calculated with Brown-Forsythe and Welch one-way ANOVA (*, P < 0.01; **, P < 0.001, ***, P < 0.0001, NS, P > 0.05). Adipocyte area was measured in 4 biological replicates with 4 images per replicate, and statistical significance was calculated with Welch’s t-test (NS, P > 0.5). Free fatty acid and glucose levels were measured in 4 biological replicates with 2 technical replicates per assay, statistical significance was calculated with Brown-Forsythe and Welch one-way ANOVA (*, P < 0.01; ***, P < 0.001, ****, P < 0.0001, NS, P > 0.05). All statistical analysis was performed using Graphpad Prism5 software.

## Author Contributions

Concept and Design: A.E.Eades, M.M.Payne, S.H.Bossmann, M.N.VanSaun

Development and methodology: A.E.Eades, M.M.Payne, S.H.Bossmann, M.N.VanSaun

Acquisition of data: A.E.Eades, M.M.Payne, L.Evans, Z.Clark

Analysis and interpretation of data: A.E.Eades, M.M.Payne, S.H.Bossmann, M.N.VanSaun

Writing, review, and editing of the manuscript: A.E.Eades, M.M.Payne, R.M.Walsh, B.B.Bye, Z.Clark, M.J.Rekowski, E.M.Morris, S.H.Bossmann, M.N.VanSaun

## Acknowledgements

This work was supported by grant CA231052 from the NCI to Dr. VanSaun. Research reported in this publication was further supported by the National Cancer Institute, Cancer Center Support Grant P30 CA168524 (PI: Jensen), with additional CCSG pilot funding for Dr. VanSaun. Additional support was provided by NSF EFRI CEE 2129617 to Dr. Bossmann (PI). We acknowledge support for the imaging core through Community Project Funding/Congressionally Directed Spending – Construction (CE1) CE1HS47377 (Jensen, PI; Bossmann, MPI). We acknowledge the Kansas University Medical Center Integrative Imaging Core with support received from the Kansas Intellectual and Developmental Disability Research Center (NIH U54 HD090216). We acknowledge the Kansas Center for Metabolism and Obesity Research COBRE for use of Solace Zone thermoneutral caging as well as the Promethion metabolic caging. We acknowledge support from the University of Kansas (KU) Cancer Center’s Biospecimen Repository Core Facility staff for performing histological work.

